# Malformations of the sacculus and the semicircular canals in spider morph pythons

**DOI:** 10.1101/2022.01.06.475233

**Authors:** J. Matthias Starck, Fabian Schrenk, Sofia Schröder, Michael Pees

## Abstract

Spider morph ball pythons are a frequently bred design morph with striking alterations of the skin color pattern. We created high resolution µCT-image series through the otical region of the skulls, used 3D-reconstruction software for rendering anatomical models, and compare the anatomy of the semicircular ducts, sacculus and ampullae of wildtype *Python regius* (ball python) with spider morph snakes. All spider morph snakes showed the wobble condition. We describe the inner ear structures in wild-type and spider-morph snakes and report a deviant morphology of semicircular canals, ampullae and sacculus in spider morph snakes. We also report about associated differences in the desmal skull bones of spider morph snakes. The spider morph snakes were characterized by wider semicircular canals, anatomically poorly defined ampulla, a deformed crus communis and a small sacculus, with a highly deviant x-ray morphology as compared to wildtype individuals. We observed considerable intra- and interindividual variability of these features. This deviant morphology of spider morph snakes can easily be associated with an impairment of sense of equilibrium and the observed neurological wobble condition. Limitations in sample size prevent statistical analyses, but the anatomical evidence is strong enough to support an association between the wobble condition in design bread spider morph snakes and a malformation of the inner ear structures. A link between artificially selected alterations in pattern and specific color design with neural-crest associated developmental malformations of the statoacoustic organ as known from other vertebrates is discussed.

## Introduction

*Python regius*, ball python, is a constrictor snake that reaches a maximum length of c. 180 cm and is native to West and Central Africa. Because of its moderate size and relatively ease of captive husbandry it has become a popular terrarium-housed exotic pet among reptile keepers and hobbyists (Jennifer 2014). The large demand for exotic python pets led to a breeding industry, in which a significant driver of growth is the development of novel color/ pattern strains through artificial breeding selection of gene mutations (Rose and Williams 2014; D’Cruze et al. 2020). Among breeders, spider morph pythons are valued because of their unusual pattern and color appearance. However, frequent neurological diseases (wobbler syndrome) have been associated with the inherited color pattern (Rose and Williams 2014).

Wobble syndrome is a collective term for various neurological disorders observed in a variety of animals. In spider morph ball python the wobbler condition may be expressed to different degrees causing side-to-side or twisting movements of the head, impaired locomotion, and difficulty striking or constricting prey items. A specific cause for the wobble syndrome is not known (Schrenk et al., in prep). Hypotheses have been proposed (Pees 2015), but none has been tested, yet. Because the wobble syndrome that is associated with spider morph python strains includes dysfunction of the sense of equilibrium, we assumed that it is either associated with central nervous defects, peripheral defects of the vestibular organ, or vertebrae malformations. Defects of the vestibular organ and vertebrae malformations should be detectable using µCT-imaging. Our study is descriptive and explorative in a sense that we investigate the morphology of individual spider morph pythons with an explicit anamnesis, and compare it to healthy wildtype animals. We refer to companion paper (Schrenk et al., in prep.) for details of the anamnesis, clinical examination and intra vitam diagnostics of the snakes studied here.

## Materials and Methods

We studied *Python regius* (Shaw, 1802) wildtype (N= 5) and spider morphs (N=4). The spider morphs were presented to the Clinic for Birds and Reptiles at University of Leipzig by private owners to determine the cause of apparent neurological symptoms. Two of the wildtype snakes were from preserved material at the clinic in Leipzig, they were euthanized for health reason not associated with any central nervous symptoms; three wildtype snakes were obtained from preserved material at LMU Munich; protocol notes describe them as healthy when killed for other research purpose. Clinical examination and health status including neurological symptoms in spider pythons have been described in detail in Schrenk et al. (in prep.). – Animals were euthanized by injecting ketamine hydrochloride (100 mg/kg; Ketamine 10%, Selectavet, Germany) combined with medetomidine (0.25 mg/kg; Domitor, Vetoquinol, Ismaning, Germany) im, then applying T61 (2 ml/kg, Intervet, Unterschließheim, Germany) intracardially.

Because of low sample size, a morphometric and statistical analysis of the inner ear structures is not possible without violating basic principles of comparative statistics. We therefore restrain ourself to a qualitative comparison of wildtype animals and spider morphs.

The study was conducted in accordance to the faculties ethical committee approval (Faculty of Veterinary Medicine; VMF EK 4/2021).

### µCT-imaging

Before scanning all heads were transferred to 70% and 95% ethanol, and then contrasted in 1% iodine solution (in 99.5% ethanol) for a minimum of two weeks. Heads were scanned at 3 different institutions. (1) Max Planck Institute for the Science of Human History, Department of Archaeogenetics, Jena; SkyScan 2110; resulting image width * height 1100 * 1100 pixel, 20 µm isotropic voxel size. (2) Zoological State Collection, Munich; Phoenix Nanotom (GE Sensing & Inspection Technologies) cone beam CT-scanner; 19 µm isotropic voxel size. (3) University of Vienna; SkyScan 1174 (Bruker); 23.9µm isotropic voxel size.

### 3D-Reconstructions

Series of original images were imported into ImageJ (Schneider, Rasband & Eliceiri, 2012; RRID:SCR_003070) and an image stack was created. The image stack was cropped, optimized for brightness and contrast, and saved in tif-format. The tif-stack was then imported using Drishti Import v2.7 and structures of interest (skull bones, bony labyrinth, membranous labyrinth, neuronal structures) were manually segmented in Drishti (Hu, Limaye & Lu 2020). Drishti-files were imported in Meshlab_64bit_fp v2021.10 (Cignoni et al. 2008) and rendered as 3D-model. Figures were labeled and plates were composed using Adobe® Photoshop CS2 9.0 (RRID:SCR_014199).

## Results

### Wildtype

The inner ear of healthy wildtype individuals of *Python regius* is housed in the prootic bone and supraoccipital bone (Figure 1A-C). Prootic and supraoccipital bones are partially overlain by the supratemporal, parietal and exoccipital bones. The columella auris is the single middle ear ossicle. It has a thin ligamentous connection to the quadrate bone. A tympanic membrane and an external ear opening are missing (like in all other snakes). The inner ear consists of the bony labyrinth and the membranous labyrinth. The bony labyrinth forms the semicircular canals (horizontal, anterior and posterior) with their respective ampullar recesses, the vestibulum and the lagenar recess (Figure 1D). The membranous labyrinth resides inside the bony labyrinth and forms the semicircular ducts with their respective ampullae, sacculus, utriculus, lagena and the ductus endolymphaticus (Figures 1, 2). Sensory epithelia are located in each ampulla, the sacculus, the utriculus and the lagena. Figure 1D documents reconstructed endocasts of the bony labyrinth while Figure 2 documents reconstructions of the membranous semicircular canals and the sacculus. The sacculus is large and spherical, and reaches dorsad between the horizontal semicircular canal. The ampullae are distinct structures at each semicircular canal and the utriculus is distinct at the crus communis. The lagena is at the ventro-medial side of the sacculus and not seen in the lateral and dorsal views presented in Figure 2.

**Figure 1.**
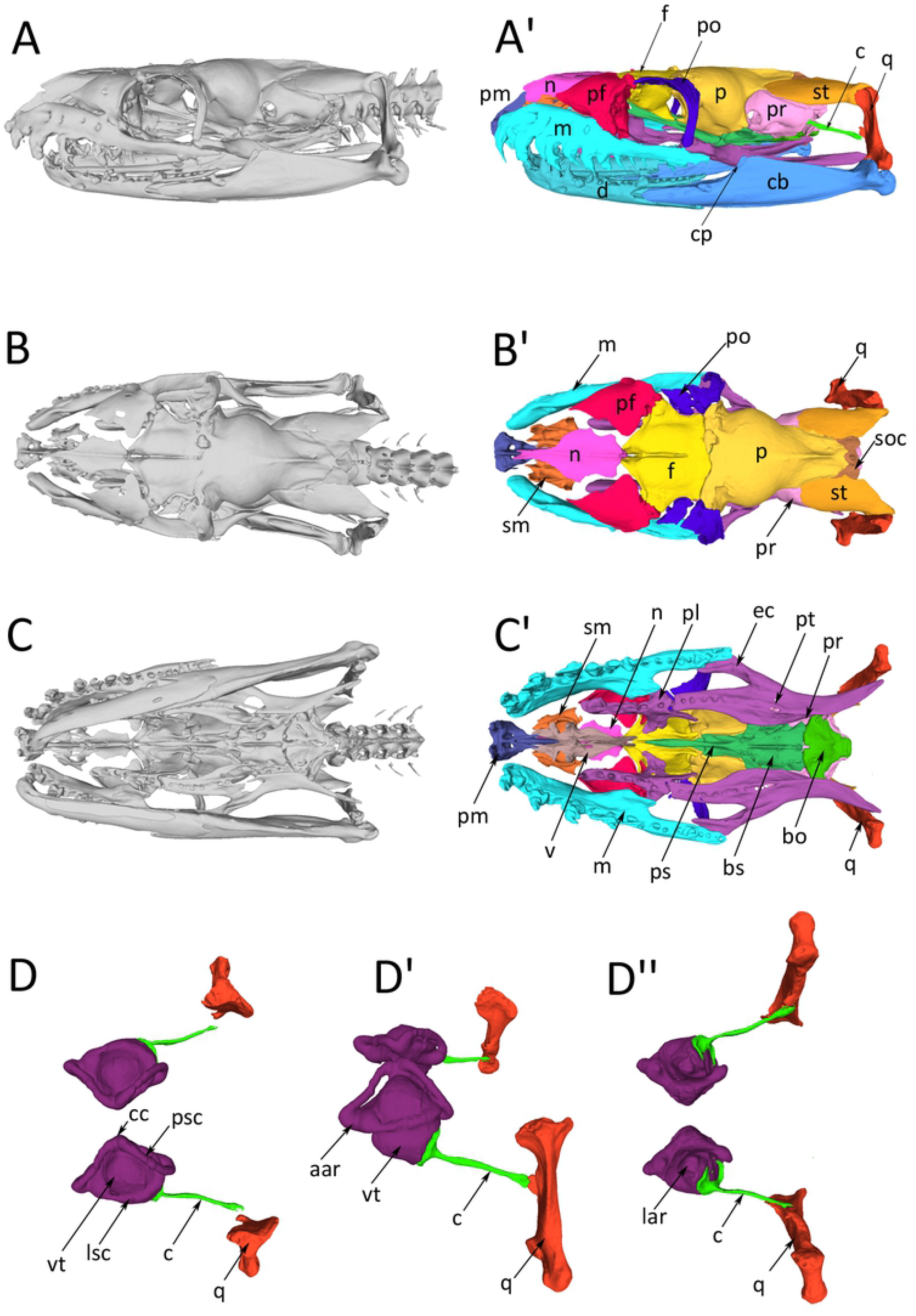
*Python regius* (# P14), reconstructions of the complete skull of a wildtype individual based on serial µCT-images. For all images left is rostral and right is occipital. The inner ear is housed in the prootic bone and supraoccipital bone. **(A)** Lateral view. (A’) Bones color coded easy distinction. **(B)** Dorsal view. (B’) Bones color coded easy distinction. **(C)** ventral view. (C’) Bones color coded easy distinction. **(D)** 3D-reconstructed endocasts of the bony inner labyrinth housing the semicircular canals and the sacculus. The columella and the quadrate are elements of the sound transmission apparatus and have been added for orientation. Dorsal (D), lateral (D’) and ventral (D’’) view. **Abbreviations:** aar, anterior ampullar recess; bo, basioccipital bone; bs, basisphenoid; c, columella; cc, crus communis; cb, compound bone; cp, coronoid process; d, dentale; ec, ectopterygoid; f, frontal bone; lar, lagenar recess; lsc, lateral semicircular canal; m, maxillare; n, nasale; p, parietal bone; pf, prefrontal bone; pl, palatinum; pm, praemaxillare; po, postorbital bone; pr, prooticum; ps, parasphenoid; psc, posterior semicircular canal; pt, pterygoid; q, quadratum; sm, septomaxilla; soc, supraoccipital bone; st, supratemporal bone; v, vomer; vt, vestibulum.

**Figure 2.**
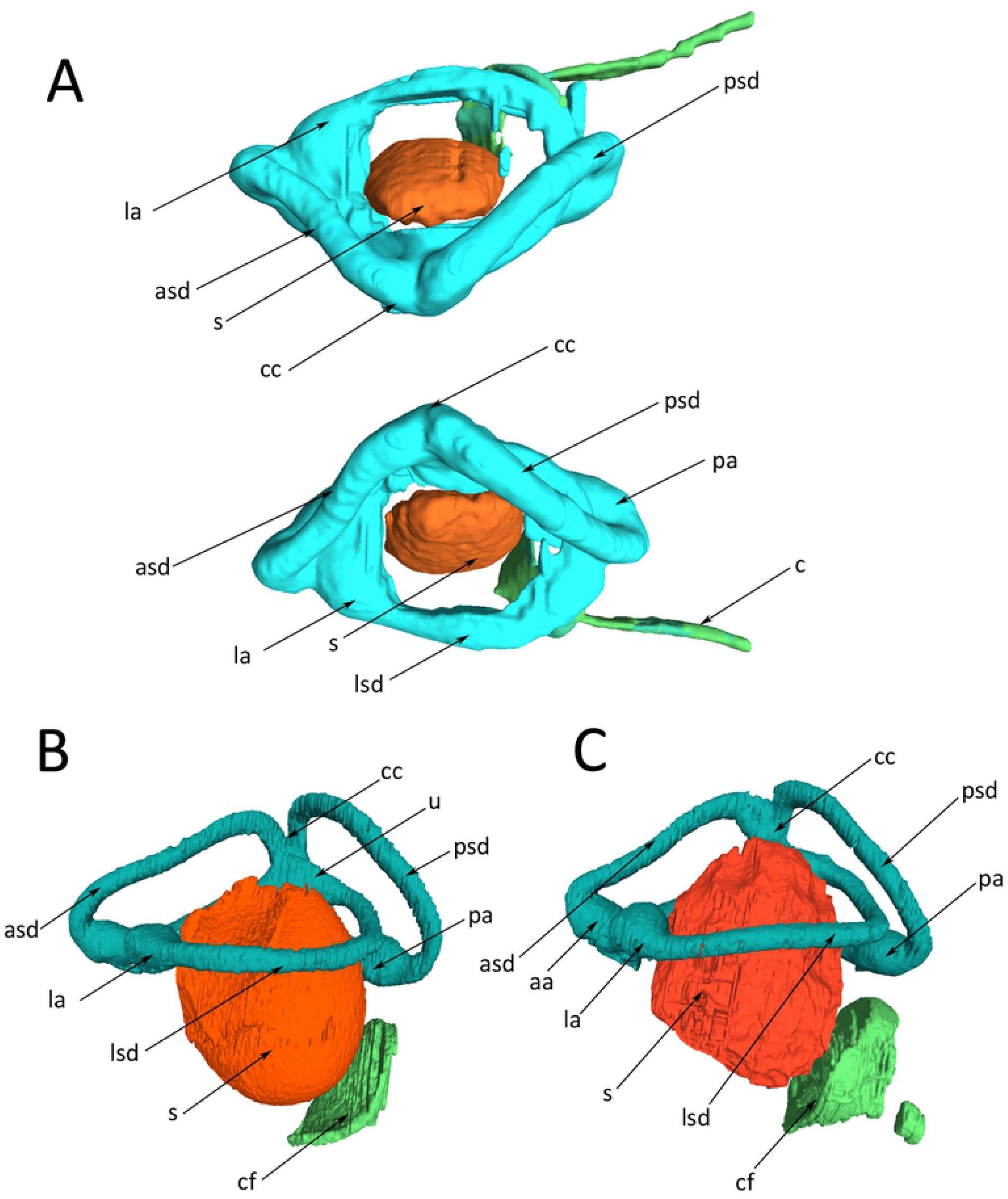
*Python regius*, 3D-reconstructions of the membranous inner ear of wildtype ball pythons. **(A)** Dorsal view of both inner ears of snake #P14. The reconstruction is based on lower resolution scans of the entire head; therefore, the reconstruction appears “cloudier” than the high-resolution scans in (B) and (C). **(B)** Right ear, control animal #1. **(C)** Right ear, control animal #2. For easy comparisons, images (B) and (C) have been mirror imaged so that left is rostral and right is occipital. **Abbreviations:** aa, anterior ampulla; asd, anterior semicircular duct; c, columella; cc, crus communis; cf, columella footplate; la, lateral ampulla; lsd, lateral semicircular canal; pa, posterior ampulla; psd, posterior semicircular duct; s, sacculus; u, utriculus.

µCT-images of the inner ear structures are sufficiently detailed for differentiation of the sensory epithelia and the membranous semicircular canals. Figure 3 documents an anterior-posterior series of selected µCT-images of the right inner ear of a healthy wildtype individual. The membranous semicircular canals are clearly recognizable in all images. They are surrounded by perilymph. In Figure 3A, the anterior ampulla and the lateral ampulla with their respective macula and the branch of the VIIIth nerve are visible. The following images show the lateral (horizontal) and the anterior semicircular canals as well as the utriculus. Figure 3C through F shows µCT-sections through the sacculus. The endolymphatic space and the macula sacculi can easily be differentiated because of their different X-ray contrast. The sacculus reaches dorsad between the lateral semicircular canal and the utriculus (Figure 3D, E). The lagena is a comparatively small extension on the medio-ventral side of the sacculus (Figure 3D, E). It cannot be seen in the lateral or dorsal views of the 3D-reconstructions (Figure 2), because it is overlain be the sacculus. – The µCT-images also provide details about the ductus endolymphaticus (Figure 3D), which, however, has not been reconstructed in Figure 2.

**Figure 3.**
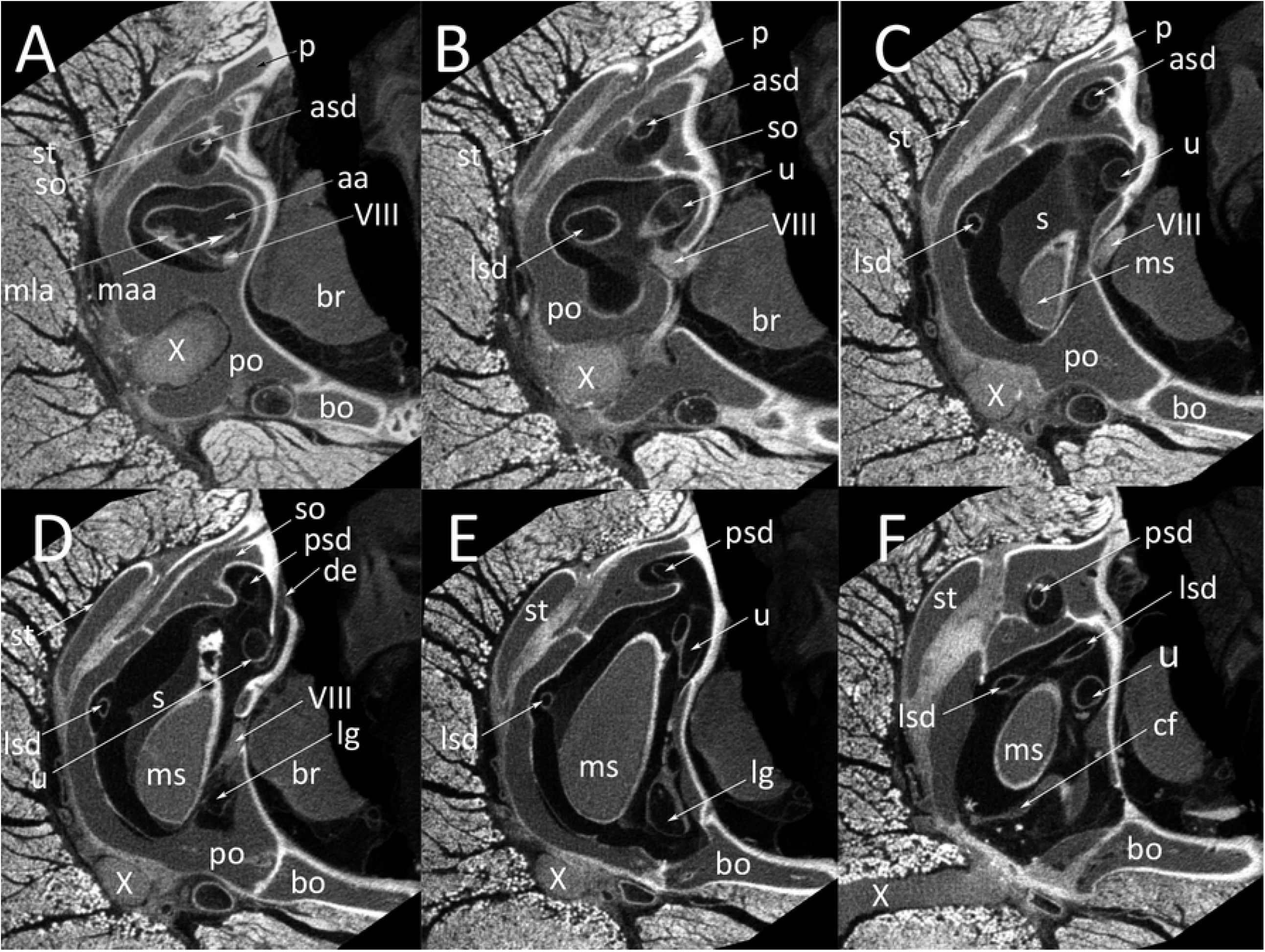
*Python regius*, serial µCT images from rostral to occipital through the inner ear in a wildtype ball python (# control animal #2). **(A)** Image #183. **(B)** Image #211. **(C)** Image #247. **(D)** Image #283. **(E)** Image #337. **(F)** Image #395. **Abbreviations:** aa, anterior ampulla; asd, anterior semicircular duct; bo, basioccipital bone; br, brain; cf, columella foodplate; de, ductus endolymphaticus; lg, lagena; lsd, lateral semicircular duct; maa, macula anterior ampulla; mla, macula lateral ampulla; ms, macula sacculi; p, parietal bone; po, prooticum; psd, posterior semicircular duct; s, sacculus; so, supraoccipital bone; st, supratemporal bone; u, utriculus; VIII, nervus stato-acusticus; X, nervus vagus.

### Spider morph

The 3D-reconstructions of spider morphs snakes render remarkably different morphology as compared to wildtype animals. The tubes of the semicircular canals are distinctly wider and the ampullae are considerably enlarged as compared to the wildtype animals (compare Figures 2 and 4). Especially the lateral and the anterior ampullae appear inflated and, because of their size merge into each other so that it becomes difficult distinguishing anterior and lateral ampullae. An external distinction is not always possible. Also, the utriculus and the crus communis are enlarged. In two individuals (Figure 4B, C) the utriculus forms a recess.

**Figure 4.**
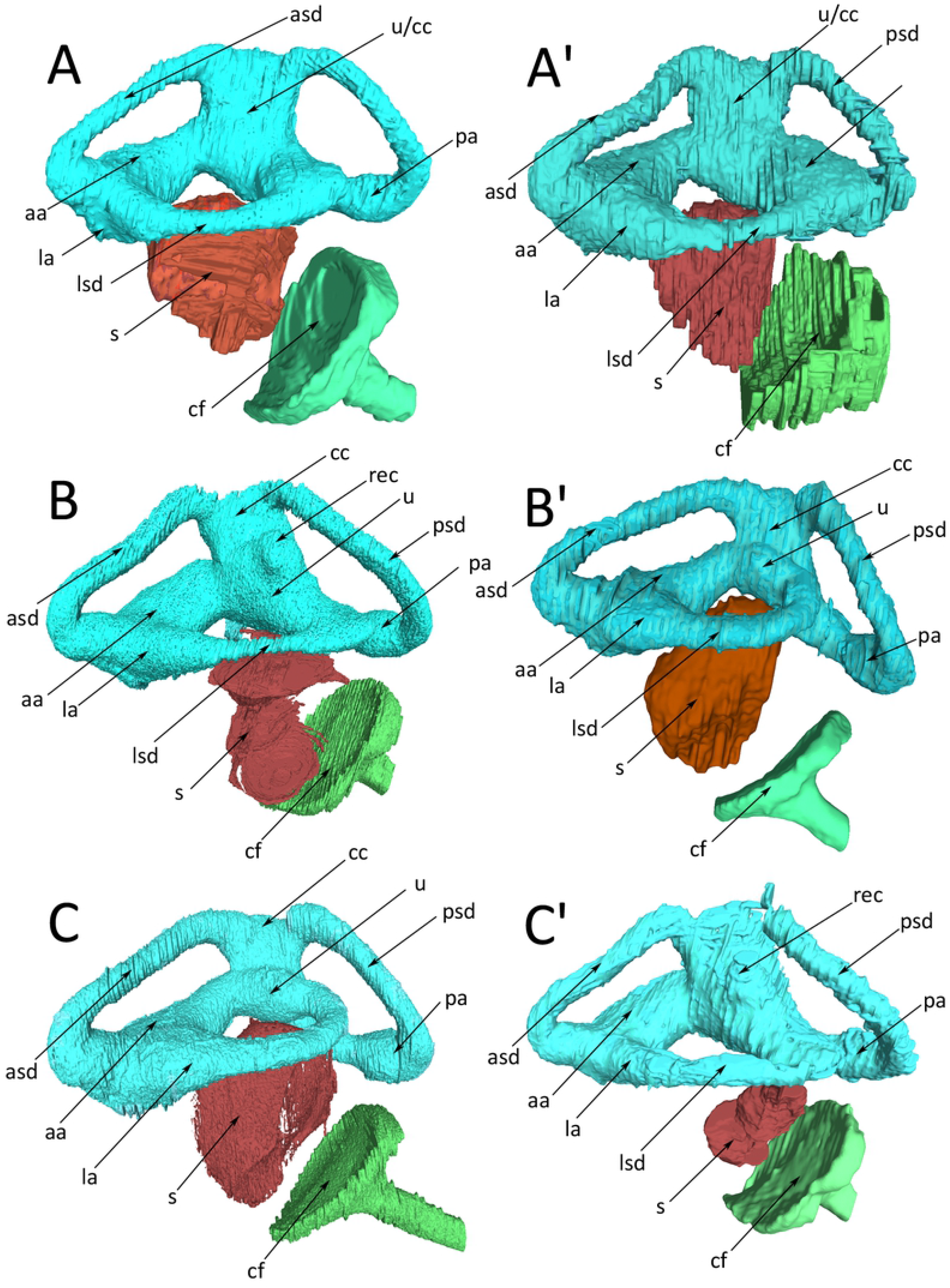
*Python regius*, 3D-reconstructions of the membranous inner ear of spider morph pythons. For easy comparisons, images (A), (B) and (C) have been mirror imaged so that left is rostral and right is occipital. **(A)** Right, **(A’)** left inner ear of snake #67532. **(B)** Right, **(B’)** left inner ear of snake #67533. **(C)** Right, **(C’)** left inner ear of snake #67894. **Abbreviations:** aa, anterior ampulla; asd, anterior semicircular duct; c, columella; cc, crus communis; cf, columella footplate; la, lateral ampulla; lsd, lateral semicircular canal; pa, posterior ampulla; psd, posterior semicircular duct; rec, recess on the utriculus; s, sacculus; u, utriculus.

The anatomy of the semicircular canals differs within individuals (left-right comparison) as well as between individuals. This is specifically evident with the lateral semicircular canal and how it connects to the anterior and posterior semicircular canals (Figure 4); e.g., in snake #67532 the lateral semicircular canal of the left side (Figure 4A’) connects to two enlarged spaces (ampullae) resulting in a largely deformed membranous labyrinth, while on the right side (Figure 4A) it looks more regular, although the ampulla of the anterior canal is also inflated. In snake #67533 (Figure 4B), the lateral semicircular canal of the left side connects directly into the utriculus, while the anterior ampulla is inflated. On the right side of this individual the lateral semicircular canals is thin in the middle part, the utriculus and crus communis are deformed. Individual #67894 (Figure 4C) has a deformed and inflated crus communis and utriculus on the left side, on the right side the lateral semicircular canal forms a small ring that connects to the anterior semicircular canal.

However, the most conspicuous difference, which was consistently found in all individuals, concerns the size and morphological integrity of the sacculus. The sacculus is distinctly smaller, never reaches through opening of the circle formed by the lateral semicircular canal and is not spherical but irregularly deformed. Left and right side of individuals may differ in sacculus structure, quite obvious when comparing Figure 4B and 4B’ or Figure 4C and 4C’.

Exemplar µCT-images document the differences to wildtype animals. Images in Figure 5 represent same sectional planes as in Figure 3, thus are directly comparable. Most conspicuous are the inflated appearance of the ampullae, the small size and diffuse morphology of the sacculus and the lack of a macula sacculi. The serial images in Figure 5 are from the left ear of snake #67533 and may be compared to the reconstructions in Figure 4B’.

**Figure 5.**
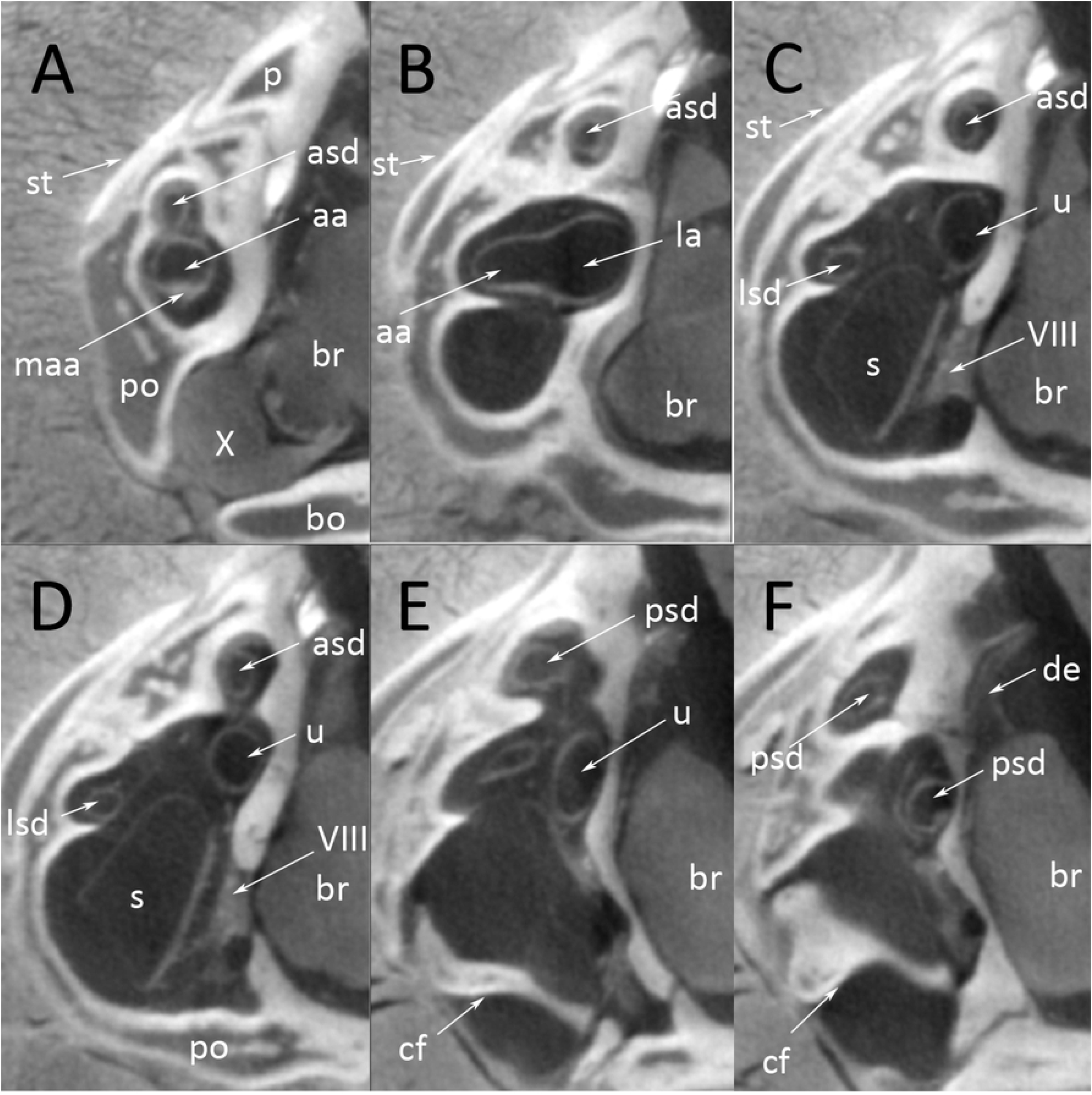
*Python regius*, serial µCT-images from rostral to occipital through the semicircular canals and the sacculus of a spider morph python (snake #67533). **(A)** Image #463. **(B)** Image # 382. **(C)** Image #355. **(D)** Image #345. **(E)** Image #275. **(F)** Image #258. **Abbreviations:** aa, anterior ampulla; asd, anterior semicircular duct; bo, basioccipital bone; br, brain; cf, columella foot plate; de, ductus endolymphaticus; la, lateral ampulla; lsd, lateral semicircular duct; maa, macula anterior ampullae; ms, macula sacculi; p, parietal bone; po, prooticum; psd, posterior semicircular duct; s, sacculus; so, supraoccipital bone; st, supratemporal bone; u, utriculus; VIII, nervus stato-acusticus; X, nervus vagus.

Intra- and interindividual differences have been highlighted above and can equally be found in µCT images of the other snakes studied. Figure 6 is a direct comparison of a cross-sectional plane through the sacculus and lagena of two wildtype snakes (Figure 6 A, B) and two spider morph snakes (Figure 6 C, D). The macula sacculi is distinct and clear in the wildtype animals. The freckling of the macula in Figure 6B is most probably caused by fixation artifact. However, the originally coherent macula structure can still be recognized. In spider morph snakes, the macula sacculi is either disorganized in a small sacculus (Figure 7C) or completely missing (Figure 7D). In spider morph snakes, the lagena is filled with a structure of low X-ray contrast. It would require histology to determine the nature of this content.

**Figure 6.**
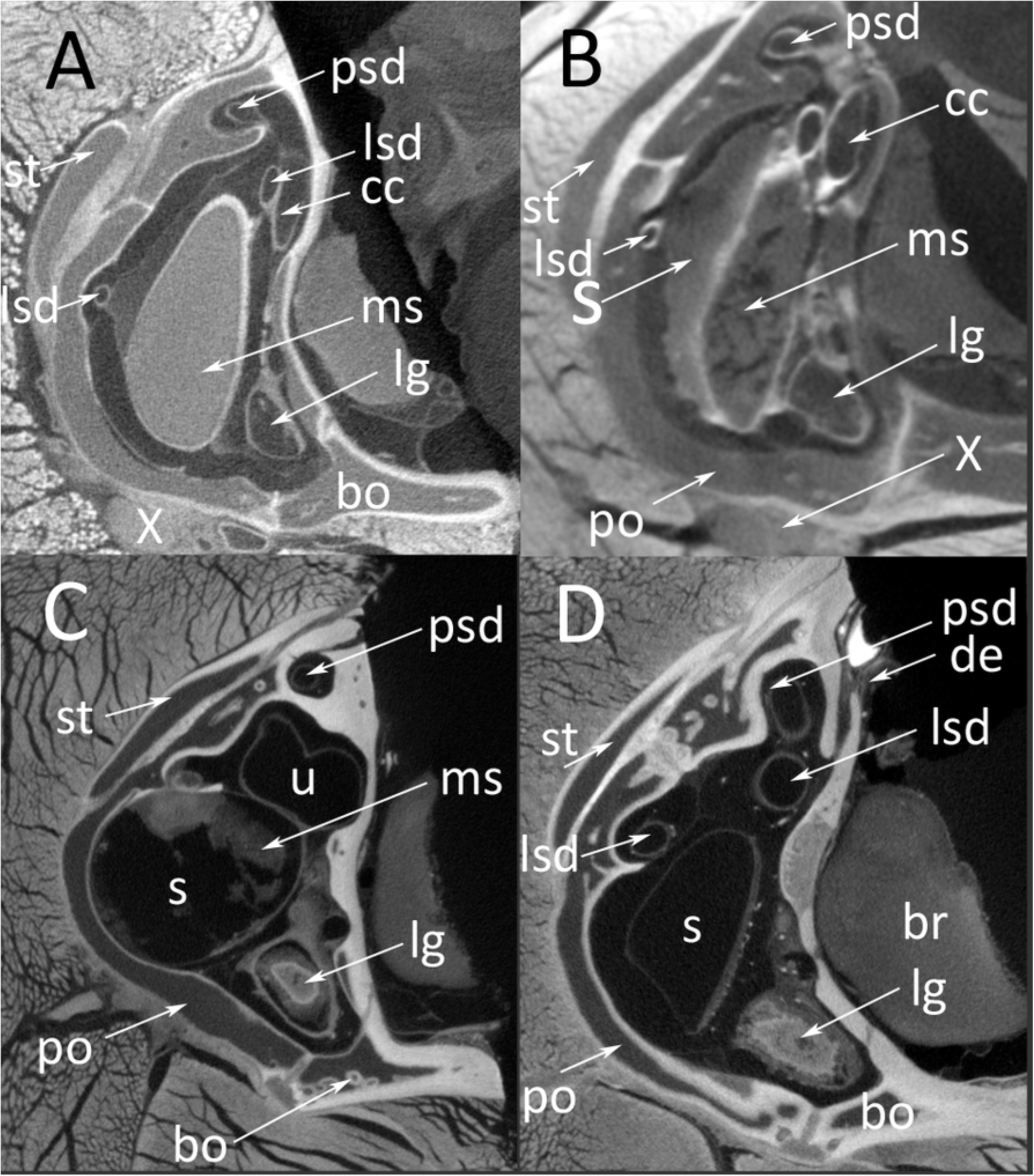
*Python regius*, µCT-images of the sacculus and lagena. **(A)** wildtype snake (control animal #1). **(B)** Wildtype snake (control animal #2). **(C)** Right inner ear of a spider morph snake (#MPI_67533/ image #511). **(D)** Right inner ear of a spider morph snake (# MPI67894 / image #628). **Abbreviations:** bo, basioccipital bone; br, brain; cc, crus communis; lg, lagena; lsd, lateral semicircular duct; ms, macula sacculi; po, prooticum; psd, posterior semicircular duct; s, sacculus; st, supratemporal bone; u, utriculus; X, nervus vagus.

**Figure 7.**
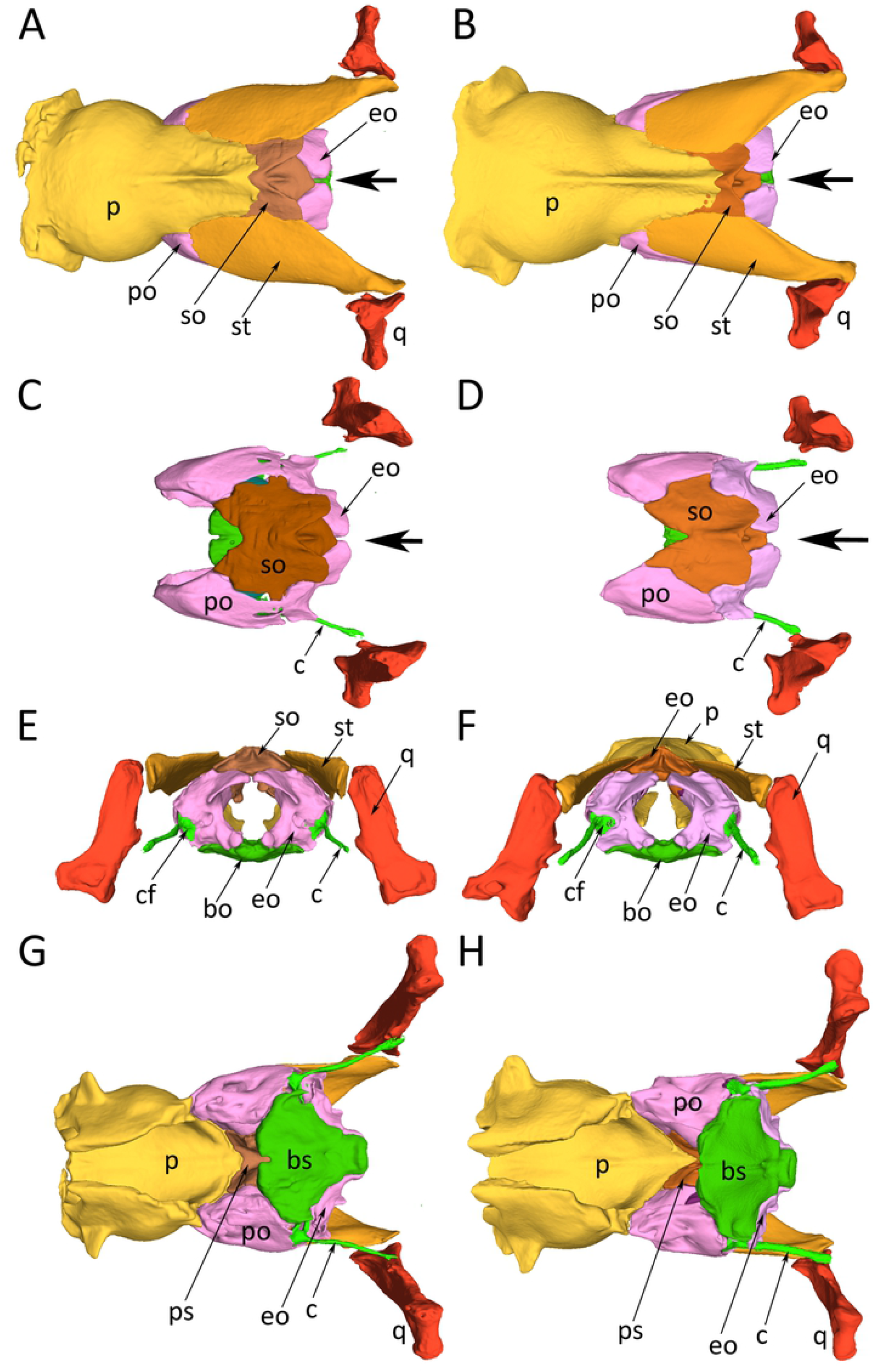
*Python regius*, reconstructions of the occipital region of skulls of a wildtype individual (left image column, snake P14) and a spider morph python (right image column, snake #70496) based on serial µCT-images. **(A)** Dorsal view on the occipital region of the skull of a wildtype snake. **(B)** Dorsal view on the occipital region of the skull of a spider morph snake. **(C)** Dorsal view on the occipital region of the skull of a wildtype snake, parietal and supratemporal bones removed so that supratemporal bones and supraoccipital bones are exposed. **(D)** Same view as (C) on a spider morph snake. **(E)** Occipital view on the skull of a wildtype snake. **(F)** Same view as (E) on a spider morph snake. **(G)** Ventral view on the skull of a wildtype snake. **(H)** Same view a in (G) on a spider morph snake. **Abbreviations**: bo, basioccipital bone; bs, basisphenoidal bone; c, columella; cf, columella foot plate; eo, exoccipital bone; p, parietal bone; po, prootic bone; ps, parasphenoidal bone; q, quadrate; so, supraoccipital bone; st, supratemporal bone; the large black arrow points to differences int eh gap between contralateral exoccipital bones.

### Desmal skull bones

The four spider morph snakes studied have a distinct morphology of the semicircular canals and the sacculus. We expected finding associated differences in the desmal skull bones. However, due to material limitations, we could compare only one pair of wildtype and spider morph snakes. We found minor differences in the size and morphology of the supraoccipital bone and the exoccipital bones. In the spider morph snake, the exoccipitals gaped distinctly wider than in wildtype snake (Figure 7 A-F). The crest of the supraoccipital was wider and more robust in wildtype snakes as compared to spider morphs. We could not find any distinct morphological differences on the ventral side of the skull. (Figure 7G, H). While these findings in a paired comparison of individual snakes have only the value of a case study, we consider them indicative of true differences between groups (wildtype vs. spider morph) because all other skull bones converge on the same morphology and do not show such remarkable differences.

## Discussion

We compare wildtype and spider morph individuals of *Python regius*. All wildtype individuals were healthy and showed no signs of wobbler syndrome, while all spider morph individuals showed noticeable neurological problems that were described as “wobble syndrome” (Schrenk et al. in prep.). Sample size in this study was primarily limited by access to material. Breeders are usually reluctant presenting their animals to the clinic because they are concerned about health issues that would prevent them from breeding. Over a period of 2 years, only 5 spider morph pythons were presented to the clinic. Sample size in this study is also limited by the time-consuming procedure of 3D-reconstruction which requires manual segmentation of all structures of interest.

µCT-imaging, although through different devices and of different contrast quality, renders cross-sectional images that are sufficiently detailed for a 3D-reconstruction of the membranous semicircular canals, ampullae and sacculus, sensory structures (maculae) as well as the bones housing the inner and middle ear structures. No published reference images of the membranous labyrinth exist besides Miller (1968), who presented a remarkable overview on the ophidian membranous labyrinth (but not for *Python regius*). However, his data were based on macroscopic dissections and are difficult to compare with the µCT-imaging presented here. All recently published papers using µCT-imaging report on the bony labyrinth (Olori et al. 2010; Boisl et al. 2011; Yi and Norell, 2015; Palci et al. 2017). A study by Christensen et al. (2012) focused on the hearing capabilities in healthy *Python regius*. This study presented also a 3D-reconstruction of the bony labyrinth and vestibule (including the macula sacculi), columella and quadrate. All published µCT data is fully consistent with our reconstructions. Therefore, we consider this independent support that methods and documentation used here are fully appropriate to describe and record differences in morphology between wildtype animals and spider morph snakes. Our data compare with classical as well as modern (µCT of the bony labyrinth) documentations of the ophidian inner ear structures.

Although we are limited in sample size, we find distinct morphological differences when comparing healthy wildtype snakes and spider morph snakes. In spider morphs, the cross-sectional diameter of the semicircular canals appears wider and the ampullae and the crus communis are inflated. The topography of the semicircular canals varies (intra- and interindividual) in the points of fusion and the width of the circle. Besides the structural differences and despite the small sample size, the inter- and intraindividual variability appears higher in spider morph snakes than in wildtype snakes.

Most conspicuous is the morphological difference of the sacculus when comparing wildtype and spider morphs. As reported above, the sacculus of spider morph snakes is distinctly smaller, deformed and lacks a coherent macula. Because the sacculus and the ampullae of the semicircular canals are organs of equilibrium, it is not surprising that these morphological differences are associated with neurological symptoms ranging from head tilt, tremor, corkscrewing, reduced striking accuracy to reduced righting reflex (reported in detail in Schrenk et al. in prep.).

### Spider morph python and vestibular disorder

Given the small sample size in our study the results rather have the character of an “extended case study” and certainly cannot be tested statistically. Therefore, conclusions should be drawn carefully. Concerns may be raised that the sample of spider snakes presented to the clinic might be biased because only individuals with wobble syndrome were presented while spider morphs without or only mild symptoms were (naturally) not presented. This may be true, but there are several lines of independent evidence that suggest an association of python spider morph strain with the occurrence of vestibular disorders.

An independent study by Rose and Williams (2014) showed that all spider morph pythons were affected by the wobbler condition, however, to some variable degree.

An association between oto-vestibular dysfunction and albinism has been reported for many albinotic animals including man and arguments for a common causative aetiology related to melanocyte function were mentioned in most of the literature (Schrott and Spoendlin 1987; Smith et al. 2000; Dawoud et al. 2017; de Jong et al. 2017). This genetic association is known since a long time and has resulted in appropriate warning against use of albinotic animals as models in biomedical research (e.g., Creel, 1980;). Reissmann and Ludwig (2000) reviewed pleiotropic effects of color-associated mutations in mammals and man. They highlight that the sensory organs and nerves are particularly affected by disorders because of the shared origin of melanocytes and neurocytes from the neural crest. Although the inner ear mainly derives from otic placode material and occipital bones housing the inner ear structures derive from axial mesoderm, an intimate interaction between migrating neural crest cells and otic neurogenesis has been described (for mammals). Neuroblasts are specified from otic vesicle epithelium, but the ganglion develops in close association with neural crest cells, which give rise to glia (Whitfield 2015).

Our study is on an exotic pet animal that has not been considered in developmental studies. Even normal healthy hearing function has been studied only once (Christensen et al. 2012). The difficulty of obtaining material and the reluctant cooperation with snake breeders prevents us from providing statistical evidence for an association of spider morph breeds and neurological disorder [but see Rose and Williams (2014) for an evidence based approach]. However, we provide explicit morphological evidence that four individual spider morph snakes have a morphology of the semicircular canals and the sacculus that is deviant from wildtype. The deformations of the sacculus, the (partial) lack of the macula sacculi, and inflated morphology of the semicircular canals suggest that the sense of equilibrium does not function properly in individuals with such deviant inner ear morphology. The conclusion is straightforward that the observed wobble syndrome in those four snakes is based on the deviant morphology of their inner ear. The intra- and interindividual variability of the deviant morphology might be related to the described variability in the expression of the wobbler condition (from light to severe). Only more exhaustive, non-invasive investigations of the stato-acoustic system in spider morph pythons can provide a sufficiently large sample size to support an association of the deviant inner ear morphology of spider morph snakes with the wobbler syndrome.

## Conclusions

We report a deviant morphology of the inner ear in four individual spider morph pythons, which all showed the wobble condition. The deviant morphology is suggestive of massively affecting the statoacoustic sense. Together with previously published evidence that all spider morph pythons are, to a different degree, affected by the wobbler condition, and together with the long-known association of color-mutants with otoacoustic diseases, we suggest that breeding for alterations in pattern and specific color design like spider morph pythons might be linked to neural-crest associated developmental malformations of the statoacoustic organ. The observed intra- and interindividual variation might account for the variability in severity of the wobbler condition. Of course, it will require more systematic investigations and a larger sample size to provide evidence of a statistical correlation between clinical symptoms of wobble disease and the extent of vestibular disorder.

## Acknowledgements

We thank Roland Melzer, Zoological State Collection Munich, Brian Metscher, Theoretical Biology Unit, Department of Evolutionary Biology, University of Vienna, and Alexander Stoessel, Max-Planck Institute for the Science of Human History, for scanning python heads. We thank Sabine Sass for her technical help in the lab.

## Author Contributions (CRediT)

**J. Matthias Starck:** Conceptualization (lead); Resources (supporting); Project administration (supporting); investigation (lead); supervision; writing – original draft (lead); formal analysis (lead); visualization (lead); writing – review and editing (lead)

**Fabian Schrenk:** Project administration (lead); Resources (lead); writing – review and editing

**Sophia Schröder:** investigation (3D reconstruction skull; desmal bones)

**Michael Pees:** Conceptualization (lead); Resources (supporting); Project administration (supporting); investigation (supporting); supervision; writing – review and editing.

## Competing interests

The authors have no competing interests to declare that are relevant to the content of this article. No funding was received for conducting this study.

## Ethical approval

The study was conducted in accordance to the faculties ethical committee approval, University of Leipzig, Veterinary Department (VMF EK 4/2021).

## Notes

### Competing Interest Statement

The authors have declared no competing interest.

